# Drift in Individual Behavioral Phenotype as a Strategy for Unpredictable Worlds

**DOI:** 10.1101/2024.09.05.611301

**Authors:** Ryan Maloney, Athena Ye, Sam-Keny Saint-Pre, Tom Alisch, David Zimmerman, Nicole Pittoors, Benjamin L. de Bivort

## Abstract

Individuals, even with matched genetics and environment, show substantial phenotypic variability. This variability may be part of a bet-hedging strategy, where populations express a range of phenotypes to ensure survival in unpredictable environments. In addition to phenotypic variability between individuals (“bet-hedging”), individuals also show variability in their behavioral phenotype across time, even absent obvious external cues. There are few evolutionary theories that explain random shifts in phenotype across an animal’s life, which we term phenotypic drift. We use individuality in locomotor handedness in *Drosophila melanogaster* to characterize both bet-hedging and drift. We use a continuous circling assay to show that handedness spontaneously changes over timescales ranging from seconds to the lifespan of a fly. We compare the amount of behavioral drift and bet-hedging across a number of different fly strains and show independent strain-specific differences in bet-hedging and behavioral drift. We show manipulation of serotonin changes the rate of behavioral drift, indicating a potential circuit substrate controlling behavioral drift. We then develop a theoretical framework for assessing the adaptive value of phenotypic drift, demonstrating that drift may be adaptive for populations subject to selection pressures that fluctuate on timescales similar to the lifespan of an animal. We apply our model to real-world environmental signals and find patterns of fluctuations that favor random drift in behavioral phenotype, suggesting that drift may be adaptive under some real-world conditions. These results demonstrate that behavioral drift plays a role in driving variability in a population and may serve an adaptive role distinct from population level bet-hedging.

**Significance Statement:** Why do individuals animals spontaneously change their preferences over time? While stable idiosyncratic behavioral preferences have been proposed to help species survive unpredictable environments as part of a bet-hedging strategy, the role of intraindividual shifts in preferences is unclear. Using *Drosophila melanogaster*, we show the stability of individual preferences is influenced by genetic background and neuromodulation, and is therefore a regulated phenomenon. We use theoretical modeling to show that shifts in preferences may be adaptive to environments that change within an individual’s lifespan, including many real-world patterns of environmental fluctuations. Together, this work suggests that the stability of individual preferences may affect the survival of species in unpredictable worlds — understanding that may be increasingly important in the face of anthropogenic change.

---

No two organisms of the same species, even when genetically identical, behave precisely the same. Individuality has been observed and measured in organisms ranging from bacteria(1) to plants(2), flies(3–7) to humans(8), even in the absence of genetic or environmental differences. This variability poses two major questions in biology—how does it arise and what, if any, evolutionary role does it serve.

Heritable phenotypic variation in a population allows for adaptive tracking, i.e., when alleles change in frequency as the selective pressure of the environment changes (9). A complementary strategy for fluctuating environments is phenotypic plasticity, in which organisms change their behavior (or morphology) in direct response to environmental changes. These two adaptive strategies share a basic outcome: the phenotype they produce matches the current or recent environment. Organisms that fail to match the environment can suffer deadly consequences.

Random differences in phenotype may reflect a “bet-hedging” strategy for species to deal with unpredictability in their environment (2, 10–16). Under this theory, variability allows some individuals to survive no matter what future environment arrives, increasing the odds that a population avoids extinction. Thus, bet-hedging species accept a lower arithmetic mean fitness for a higher geometric mean fitness (which equals zero if there is a single generation of no fitness). Theoretical work shows that bet-hedging strategies relying on random non-heritable variation outperform adaptive tracking in some environments (6, 16, 17). In particular, bet-hedging outperforms adaptive tracking when the environment fluctuates on timescales similar to the lifespan, as adaptive tracking requires multiple generations to respond to environmental cues (15, 18), and can lag behind fluctuating selective pressures. While phenotypic plasticity occurs on faster timescales, the ability to adapt and learn can be metabolically costly and provides limited buffer against sudden changes in selective pressures (19). Intrinsic variability in genetic control (1) has been identified as a source of phenotypic variation in microbes. In multicellular organisms, stochastic processes in development underlie variation (20), and, in the case of behavior, stochastic neuronal wiring (4, 21–23) has been shown to predict persistent individual differences.

Individuals, however, do not express a singular behavioral phenotype across their life. Individual biases in multiple behavioral settings are only partially consistent over time (3, 5, 24–27), showing spontaneous changes even in the absence of macroscopic cues that could trigger plasticity. These observations raise two questions: to what degree does individuality arise due to developmental differences versus spontaneous fluctuations within the lifetime of the animal, and do fluctuations across the lifetime of an animal provide an evolutionary benefit? To answer these questions, we used near-isogenic animals to measure 1) the extent of variation present at the start of adulthood (“bet-hedging”) and 2) the amount behavioral phenotypic change over time (“behavioral drift”). We use locomotor handedness in *Drosophila melanogaster* as a model to measure how behavior drifts over time and show that the extent of behavioral drift is influenced by genes and the neuromodulator serotonin. We then use a life-history model to test the hypothesis that random change in phenotype (“phenotypic drift”, as a more general form of behavioral drift) can be adaptive and assess whether real-world environmental fluctuations may drive the evolution of phenotypic drift. Taken together, these experiments and analyses suggest that phenotypic drift is plausibly an adaptive strategy to cope with environmental fluctuations within an organism’s lifespan.

## Results

### Individual behavioral biases drift over time

To investigate the timescales on which individual preference spontaneously changes, we placed 252 flies in individual circular arenas and continuously tracked them for up to 30 days. At each time point, we computed the direction of circling (Figure 1A), a measure that exhibits idiosyncratic variation, when averaged over long timescales (3). To look for slow changes in bias, we examined the turning bias of individual flies at different timescales: first, we averaged their direction of circling over each hour and second we low-pass filtered their continuous turning behavior to discard frequency components faster than 24 hrs (Figure 1B). Comparing both of these measures to the experiment-wide average preference for each fly shows substantial long-duration changes from their experiment-wide circling tendency. To quantify this across all flies we calculated the average power spectrum of the raw turning data across flies (Figure 1C), showing substantial power in lower frequencies corresponding to hours-to days-long fluctuations in turning preferences. Interestingly, we did not see peaks at any specific frequencies (e.g., circadian), though there was a broad shoulder of power between 10^−4^ and 10^−2^ Hz. The overall trend across six orders of magnitude bore some resemblance to a power-law relationship between frequency and power. We saw similar patterns in other measures of behavior from this experiment, including speed, heading velocity, and distance from the center of the arena (Fig S1A-D).

**Fig. 1.**
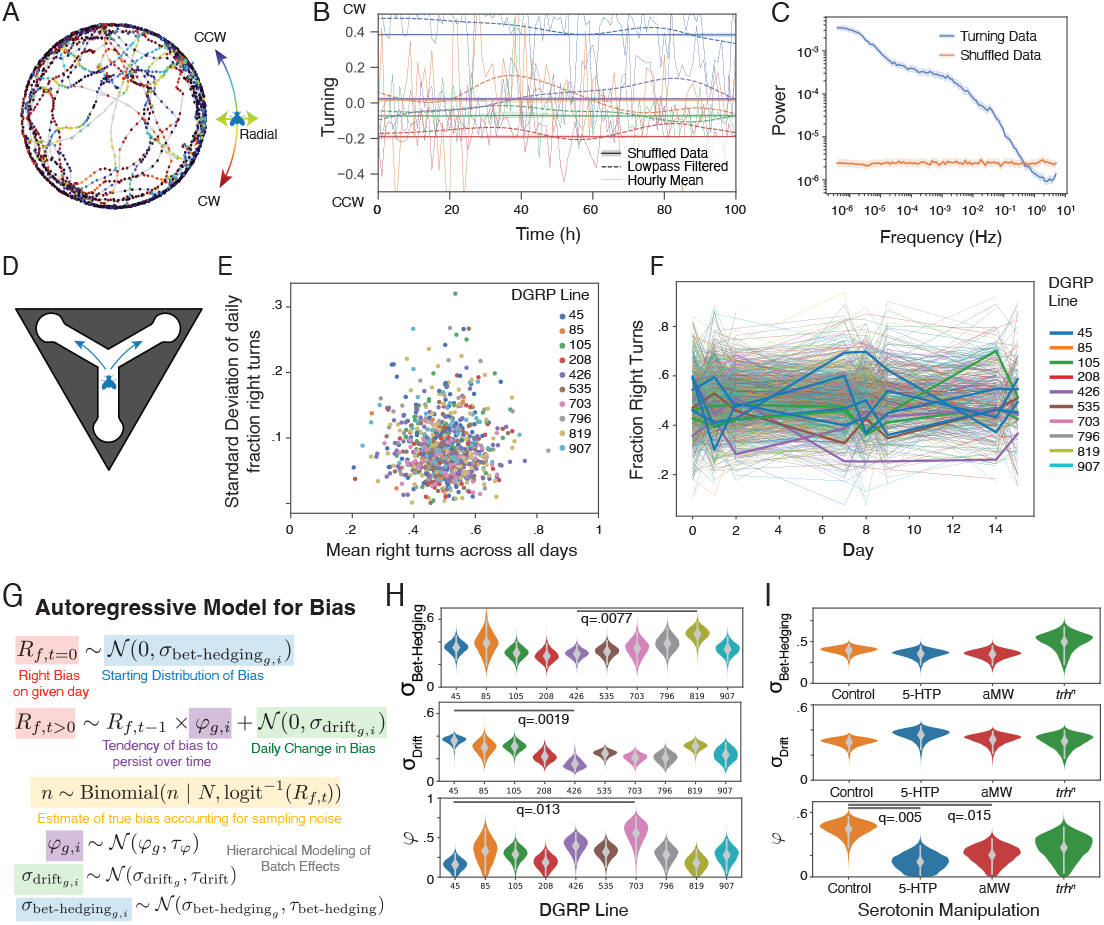
Characterizing changes in individual preferences in *Drosophila melanogaster*. **A**. 2 hr sample of centroid-tracking data for a fly in a circular arena. Each point is colored based on whether it is moving CCW, CW, or radially in the arena. **B**. Sample of 100h of continuous recording for 4 individual flies (colors). Hourly means of turning indices are shown in light lines. Dashed lines are low-pass filtered with a timescale cutoff of 24 h. Solid lines are low-pass filtered shuffled data showing the average tendency across the experiment. **C**. Mean power spectrum of turning data for all continuously monitored flies (n=252) for actual and shuffled data. Shaded areas represent 95% confidence intervals generated via bootstrapping (n=1000). **D**. Schematic of Y-maze assay. Flies make either a left or right turn each time they walk through the intersection. **E**. Standard deviation of daily right biases vs average right bias across days for individual flies (points). Colors indicate DGRP genotype (*n*=48-235 for each genotype (see Table S1)). **F**. Mean individual handedness per day across all DGRP lines, a random subset are thickened to show representative changes in individual handedness over time. **G**. Autoregressive model of individual right bias over time with parameters to estimate the initial right bias variability (*σ*_Bet-Hedging_) and rate of daily change in right bias (*σ*_Drift_). **H**. Posterior estimates of *σ*_Drift_, *σ*_Bet-Hedging_, and *φ* (the autoregressive parameter characterizing the rate of reversion to zero bias), for each DGRP genotype. Grey bars represent 95% credible intervals. Lines indicate the smallest q value between populations. **I**. As in H, posterior estimates of right bias variability parameters for flies treated with 5-HTP, AMW, and controls, as well as mutant flies with a missense mutation in *trh* generated by in vivo CRISPR. *n*=98 or 192 for each condition (see Table S1). q values *<*.05 are indicated.

### Genes regulate the rate of behavioral drift

The extent of behavioral variability in a population differs between genotypes (28–30). We next looked to see if the rate of behavioral drift also differs between genotypes. This is a prerequisite if behavioral drift can evolve as a trait under natural selection. We used 10 different lines from the *Drosophila* Genetic Resource Panel (DGRP) (31). Using a Y-maze assay (Figure 1D) that measures left-right choices that correlate with locomotor handedness in circling(3), we measured the locomotor handedness of flies in each of these lines three times weekly for three weeks. As in the circling assay (Figure 1A-C), individual flies exhibited substantial variation in their daily average turn biases (consistent with drifting biases) in all genotypes, as measured by the standard deviation in the daily fraction of right turns for each fly, showing substantial variation from their experiment-wise average (Figure 1E), and evident in plotting the fraction right turns for each day of all flies (Figure 1F). We used a hierarchical Bayesian autoregressive model (Figure 1G) to estimate the initial variability in turn bias for each genotype (*σ*_Bet-Hedging_), the extent to which turn bias changed each day (*σ*_Drift_), and the tendency of turn bias to persist over time (*φ*) (Figure 1G). We observed large differences in all three parameters between genotypes, as evidenced by minimally overlapping posterior distributions of estimates of these parameters (Figure 1H). All genotypes showed *φ* above zero, demonstrating some level of persistence in individual biases over time, consistent with our circling experiments and previous studies of Y-maze handedness (3). Interestingly, we saw independent variation across genotypes in the posteriors for *σ*_Bet-Hedging_ and *σ*_Drift_, suggesting that different genetic mechanisms regulate behavioral bet-hedging and drift.

### Manipulating serotonin modulates the rate of behavioral drift

Neuromodulators have been shown to play an important role in regulating individuality in a large range of organisms (8, 32– 35), including flies (5, 36, 37). To test whether serotonin affects the rate of behavioral drift (and/or bet-hedging), we fed adult, isogenized Oregon-R flies food supplemented with either the serotonin synthesis inhibitor aMW or the serotonin precursor 5HTP and measured their locomotor handedness for 3 days each week for 3 weeks, as in the previous experiment. While *σ*_Bet-Hedging_ and *σ*_Drift_ were similar in all three treatment groups, both pharmacological manipulations of serotonin decreased the stability of behavior over time, increasing *σ*_Drift_ and decreasing *φ* (Figure 1I, S1F). This was also evident in decreased correlation coefficients between handedness on successive days (Figure S1H-J) compared to control flies.

To assess the role of serotonin as a modulator of behavioral drift by a second approach, we generated a constitutive mutation in the tryptophan hydroxylase gene (*trh*), which is involved in the synthesis of serotonin. This approach also controls for off-target pharmacological effects and, unlike the previous experiment, deprives the animals of serotonin during development as well as after eclosion. We used *in vivo* CRISPR (38) to knock out *trh* in an isogenized Oregon R background, yielding control flies of a closely matched genetic background. We did not see clear evidence of difference from control flies in the marginal distributions of *σ*_Bet-Hedging_, *σ*_Drift_ or *φ* in our hierarchical model(Figure 1I, S1G), though we did see a significant decrease of day-to-day correlation between control and *trh* null flies (Figure S1K-M).

### A theoretical adaptive advantage for phenotypic drift

The variation we see in *σ*_Drift_ between genotypes suggests that behavioral drift can potentially evolve to provide fitness advantages in some situations. To build an intuition for the ideal phenotypic drift strategy and explore possible theoretical foundations, we formulated a simple two-state model (Figure 2A). Imagine an organism in an environment that shifts probabilistically between two states. The organism can exhibit two behavioral phenotypes. If the chosen phenotype matches the environment, the organism survives; if not, it dies. What is the optimal strategy for changing its behavior to survive probabilistic shifts in the environment that occur with probability *p*? We can prove, through methods analogous to classic findings on optimal betting solutions (39, 40) and previous analytical work on generational bet-hedging (41, 42), that in this two-state model, the optimal fraction of the population that should shift their behavioral phenotype in order to maximize long-term population growth equals *p*. This result extends to both the case of an arbitrary number of possible states as well as a continuous distribution of environmental states and behavioral phenotypes for all cases where the fitness narrowly depends on a tight match to a given environment (see Supplementary Materials for proofs). Interestingly, this holds true even when the fitness associated with successfully matching phenotype to environment A differs from matching environment B – in other words, matching the environment as much as possible is the most important thing, independent of the quality of specific environments.

**Fig. 2.**
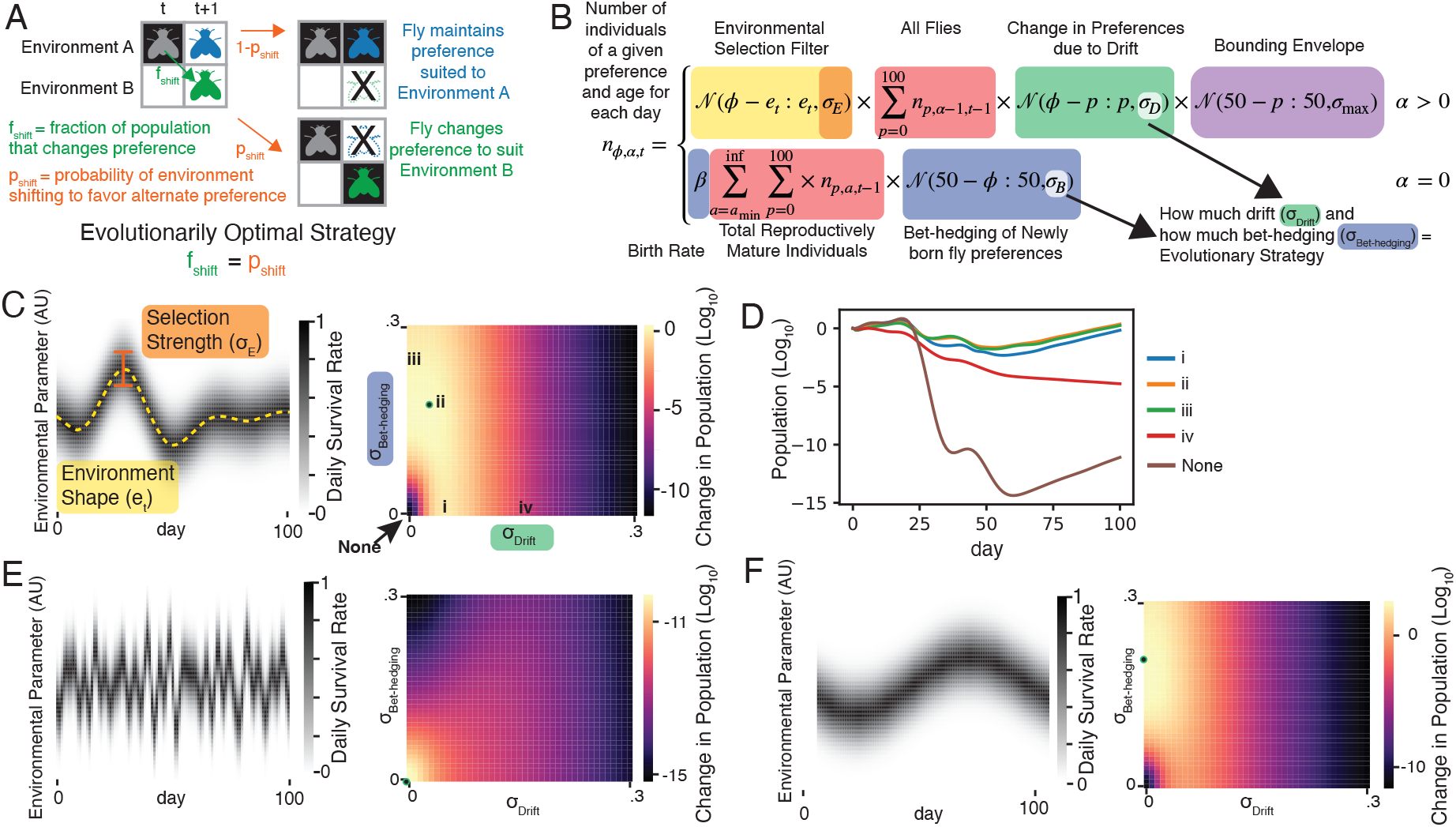
Modeling adaptive scenarios of phenotypic drift. **A**. In a simplified model in which both phenotypes and environments have two states, the optimal fraction of the population that should change preference (*f*_shift_) over a period of time equals the probability the environment changes (*p*_shift_). See Supplementary text. **B**. Model for the number of individuals with a particular continuous preference as a function of time in a fluctuating environment and individual age. *n*_*ϕ,α,t*_, is the number of individuals with preference *ϕ*, age *α* at day *t* in the simulation. Two different cases determine this value, one for flies surviving from the previous timestep (*α >* 0), and one for flies born in a particular timestep (*α* = 0). The number of flies of a particular behavioral phenotype surviving on each successive day is determined by a function of how far that preference is from the ideal preference on that day (orange), the total number of flies that already have, or drift into having that phenotype on that day (red), and a bounding term (purple) that stops the distribution of preferences from diffusing away from general range of what is adaptive (adding some degree of reversion to the mean consistent with our observed values of *ϕ*, and via the Wiener-Kninchin theorem, consistent with the power spectrum observed in Fig 1C). The key behavioral strategy parameter from these terms is *σ*_*d*_, which determines the rate at which flies’ preferences drift over time. The number of new flies born each day is given by the total number of flies above the age of reproductive maturity *a*_min_ (red) times the birth rate *β*. New flies are born with an initial preference from a normal distribution centered on the long term environmental mean with a standard deviation given by *σ*_bet-hedging_ (blue) **C**. Example environmental fluctuations and corresponding fitness landscape showing the change in population over 100 simulation days for differing amounts of phenotypic drift and bet-hedging. Green dot indicates ideal strategy (ii)**D**. Population over time for strategies marked with roman numerals in (C). **E, F**. As in (C) for two additional example environmental fluctuation patterns.

### A flexible model of phenotypic drift shows a potential adaptive role under some patterns of environmental change

To model the effect of more realistic environmental fluctuations, we added dynamic environments, fitness effects and life history to the auto-regressive model we used to fit the experimental behavioral data (Figure 2B). In this model, individuals, tracked at the population level, exhibit different behavioral preferences that can potentially vary over their lifespans. The environment fluctuates, and the survival probability of an individual increases as the difference between their preference and the environment decreases. Two parameters determine the phenotypic strategy: *σ*_Bet-Hedging_, captures the initial variability of the population, and *σ*_Drift_, captures the amount each fly’s preference changed per day. By integrating this model over time, we estimated the fold-change in population size with varying levels of *σ*_Bet-Hedging_ and *σ*_Drift_ (Figure 2C-D, S2), thereby determining the optimal variability strategy for a given pattern of environmental fluctuations. Sampling a few randomly generated environments quickly revealed that different patterns of fluctuation favored different strategies (Figure 2E-F), so we began to systematically assess what characteristics of the environment and life history might favor phenotypic drift, bet-hedging or combinations of both.

### Phenotypic drift is an effective response at shorter timescale fluctuations than bet-hedging

To characterize the adaptive value of bet-hedging and phenotypic drift strategies as a function of statistics of environmental fluctuations, we created randomized environmental dynamics by filtering white noise in the time domain, leading to random fluctuations with a given temporal frequency content. We normalized and scaled the resulting environmental patterns, and used them in simulations of population survival for a range of bet-hedging (*σ*_Bet-Hedging_) and phenotypic drift (*σ*_Drift_) values. From these simulations we constructed a fitness landscape over the four dimensions of *σ*_Bet-Hedging_, *σ*_Drift_, environmental fluctuation period and environmental fluctuation amplitude (Figure 3A). At very high frequency fluctuations the most successful strategy was *σ*_Bet-Hedging_ = *σ*_Drift_ = 0, reflecting an optimal strategy of all individuals tightly matching the average conditions of the environment (Figure 3C). Similarly, low amplitude fluctuations favored low bet-hedging and low drift. These findings make sense: low amplitude fluctuations are effectively a static environment that does not require a variable phenotypic strategy, and very rapid fluctuations changing within the timescale of organismic response are averaged away (6, 15, 18). However, as the amplitude of the fluctuations increased and the frequency decreased, the optimal strategy showed increased amounts of drift and bet-hedging, as producing individuals with a preference far from the environmental mean became increasingly necessary for survival. While both drift and bet-hedging are preferred over no variability in these situations, higher frequency fluctuations favor drift, while lower frequency fluctuations favor bet-hedging. Increasing the time to sexual maturity increases the advantage of drift, suggesting that varying a phenotype dynamically provides a mechanism for organisms to survive long enough to reproduce. These results suggest that the relative benefit of a drift or bet-hedging strategy depends on the rate of fluctuation of the environment compared to the development time of an organism. These relationships hold for a wide range of birthrates *β* (Figure S3A) and ages of reproductive maturity *a*_min_ (Figure S3B-C).

**Fig. 3.**
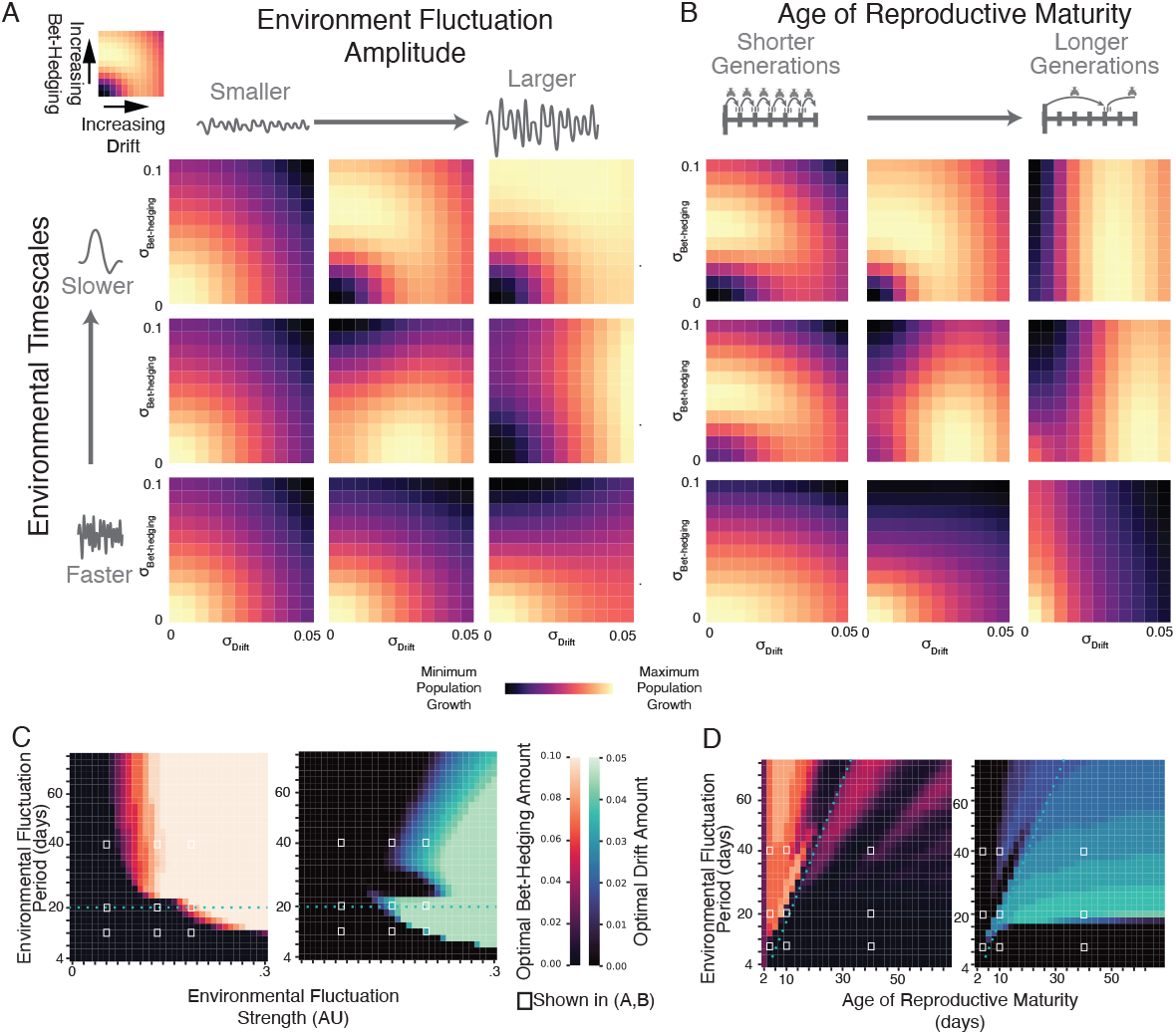
Effects of amplitude of environmental fluctuation, frequency of fluctuation, and age of reproductive maturity on ideal amounts of bet-hedging and phenotypic drift. **A**. Fitness landscapes over combinations of environmental fluctuation amplitude and frequency. Each heatmap shows the geometric mean of the log change in population for each combination of drift and bet-hedging over 100 randomized environments. Heatmaps are normalized to their maximum and minimum values. Rows of heatmaps have the same environmental fluctuation frequency and columns have the same environmental fluctuation amplitude (as measured by the standard deviation of all timepoints *σ*_Mean_. The nine amplitude and frequency combinations in this panel correspond to values denoted with white boxes in (C). **B**. As (A), except columns of heat maps have the same age of reproductive maturity, as determined by *a*_min_. **C**. Optimal amounts of bet-hedging (warm color scale; left) and drift (cool color scale; right) for each combination of environmental fluctuation amplitude and frequency. White squares indicate values associated with heatmaps in A. Dotted blue line corresponds to an environmental fluctuation period of 20 days, which is twice the age of reproductive maturity in these simulations. **D**. As (C), except for optimal amounts of bet-hedging and drift for each combination of environmental fluctuation frequency and *a*_min_. Dotted blue line corresponds to environmental fluctuations of twice the age of reproductive maturity. White squares indicate values associated with heatmaps in (B).

### Real-world environmental fluctuations may favor phenotypic drift

To predict the adaptive value of phenotypic drift strategies in somewhat less synthetic circumstances, we collected 43 types of environmental time series from publicly available National Oceanography and Atmospheric Administration (NOAA) (43) and National Ecological Observatory Network (NEON) (44) datasets across 118,618 sites. We randomly sampled contiguous 1000-day time series from each of these datasets from the past 20 years, z-scored each time series, then scaled them to have standard deviations *σ*_mean_ to represent different amplitudes of selective pressure. We then used these time series as model inputs to calculate the optimal amounts of phenotypic drift and bet-hedging as a function of *σ*_mean_ and *a*_min_ (Figure 4A-B). Different patterns of environmental fluctuation, across differing measurement types and locales, showed differing optimal amounts of phenotypic drift and bet-hedging. As in our previous analyses, increasing *σ*_*mean*_ increased the optimal *σ*_Drift_ and *σ*_Bet-Hedging_ (Figure 3), and increasing the *a*_min_ increased the optimal magnitude of phenotypic drift (Figure 4A, S4B,D).

**Fig. 4.**
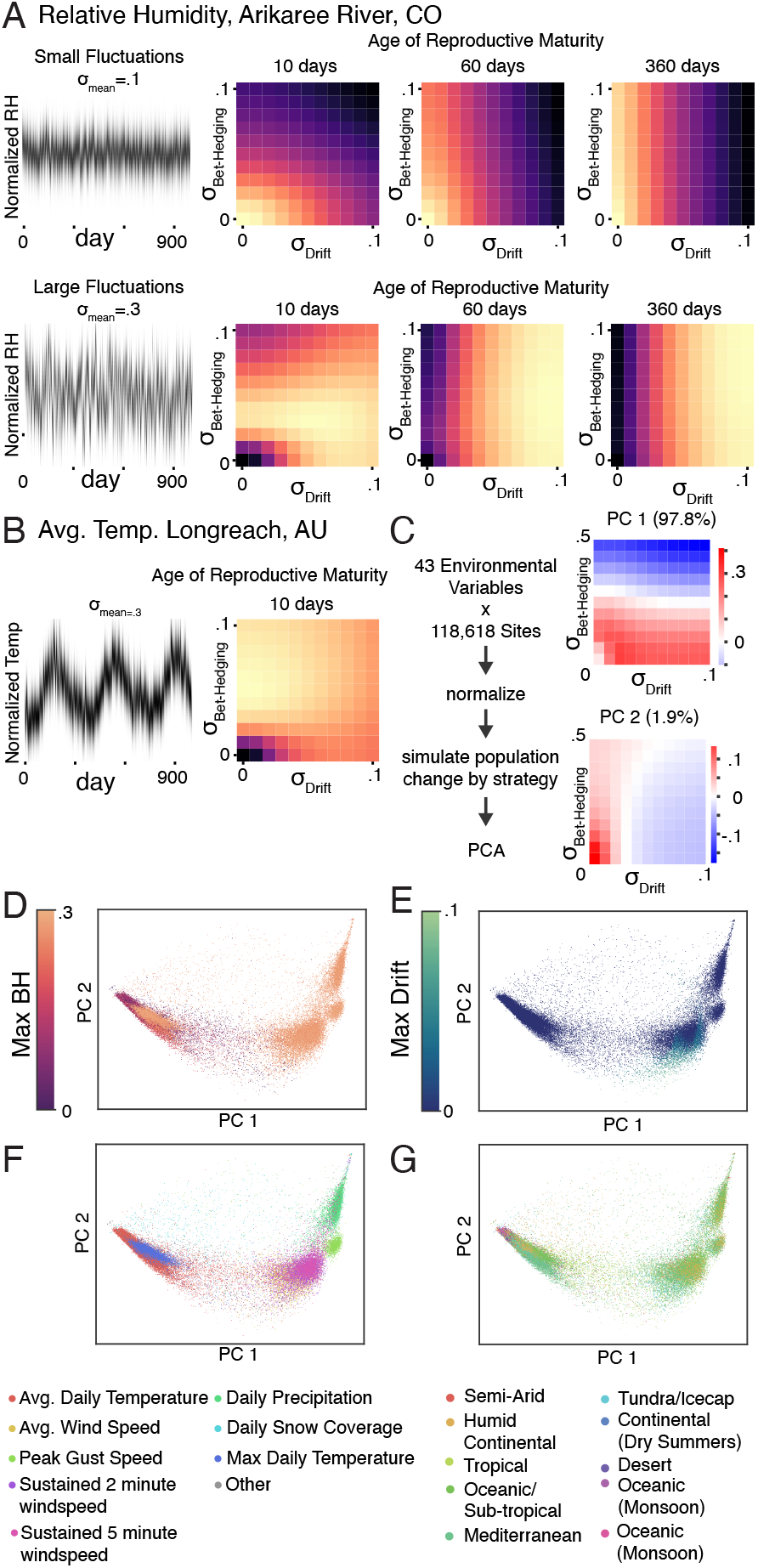
Optimal bet-hedging and phenotypic drift strategies for real-world environmental fluctuations. **A**. 1000 days of relative humidity data from the Arikaree River in Colorado, USA (left panels) were used to generate an environmental selection filter with low (top-left) or high (bottom-left) amplitude fluctuations (*σ*_mean_ in Figure 2B). Fitness landscape heatmaps over bet-hedging and drift strategies for different ages of reproductive maturity (*a*_min_). **B**. As in (A), using average daily temperature data from Longreach, Australia. **C**. Pipeline for comparing the optimal variability strategies of organisms subject to real-world environmental fluctuations. Daily environmental time series from many sites were collected, normalized, and used in the model to produce fitness landscapes over *σ*_Bet-Hedging_ and *σ*_Drift_. All landscapes were then subject to principal components analysis. These simulations held *a*_min_ and *σ*_mean_ constant. See Methods. The loadings of PC1 (97.8% of the variance; top-right) indicate that this component encodes the optimal amount of bet-hedging, while PC2 (1.9% of the variance; bottom-right) encodes optimal drift. **D**. Environmental time series from specific locations, plotted on PC2 vs PC1 axes, colored by optimal amount of bet-hedging. **E**. As in (D), except color indicates optimal amount of drift. **F**. As in (D), except color indicates the type of environmental measurement. **G**. As in (D), except color indicates the Kö ppen climate classification of their location.

To relate optimal strategies of phenotypic drift and bet-hedging to climate and environmental factors, we used principal components analysis (PCA) to study variation in fitness landscapes over *σ*_Bet-Hedging_ and *σ*_Drift_ (Figure 4C). The first principle component of variability strategy landscapes had the vast majority of the variance (97.8%) and encoded the extent of bet-hedging; the second component (1.9%) encoded the extent of drift. Across time series, we observed a variety of optimal amounts of phenotypic drift and bet-hedging. A substantial minority (12%) of environmental signals favored non-zero drift (Figure 4D, E), particularly those reflecting temperature and wind-speed measurements and a subset of locations with tropical, sub-tropical, Mediterranean and continental climates (Figure 4F, G). These results depend on the *a*_min_ in our model, with higher ages of reproductive maturity favoring more phenotypic drift (Figure S4).

## Discussion

We found that individual behavioral biases are not stable over an animal’s lifetime, and that the degree of stability in behavioral bias is influenced by genetics and neuromodulation. Inspired by these findings, we provide a theoretical rationale for how instability in preferences could be adaptive for the long-term survival of individuals and species.

The behavioral preferences of individual flies change continuously over their life on days-long timescales (Figure 1 A-C; S1A-D). While indications of this phenomenon have been seen previously (3, 5, 24, 45–47), these studies examined a small number of time points or recorded behavior continuously for up to a week. By measuring behavior continuously for up to four weeks (roughly the lifespan of a fly), we were able to estimate the complete power spectrum of behavior: the continuous distribution of timescales at which it changes. Across multiple behaviors (Figure 1C, Figure S1A-D), we show that there is no specific timescale over which behavioral preferences change (e.g., we did not see a circadian cadence to behavioral changes), and our data suggest that individuality shifts on a number of different time scales.

Examining several different inbred strains derived from a natural fly population, we found that the stability of preferences depends on genotype (Figure 1H). This implies that there is natural allelic variation that affects the rate of behavioral drift. In turn, this suggests that behavioral drift is not strongly maladaptive, in which case we would expect alleles involved in promoting behavioral drift to have been selected out of the population. Genetic variation for behavioral drift implies that the rate of drift could evolve to increase the fitness of animals subject to different kinds of environmental fluctuations. However, genetic variation for behavioral drift does not, *per se*, demonstrate that drift is subject to selection; drift could be a neutral epiphenomenon of some other trait under selection.

Importantly, some lines with differing magnitudes of initial preference variability (*σ*_bet-hedging_) exhibited similar amounts of preference stability (*σ*_drift_) (e.g., lines DGRP 819 and 426). This suggests that mechanisms that drive differences in population preferences developmentally incompletely overlap with mechanisms that drive stability in preferences as adults.

Serotonin has been implicated as a regulator of the extent of individuality in fly behavior (5, 36, 37) and stability of idiosyncratic preferences in *C. elegans* (33, 48). Bidirectional pharmacological perturbation of serotonin signaling (increasing it with serotonin precursors, or decreasing it with synthesis inhibitors or genetic perturbations) decreases the stability of preferences over time (or conversely increases the amount of behavioral drift) (Figures 1I, S1F-M). Our results from constitutive mutations in the serotonin synthesis pathway are less compelling (though we still see a significant difference in the strength of day-to-day correlations). The more modest impact of constitutive *trh*^*n*^ mutation may reflect compensatory effects playing out across development. Together, our serotonin manipulation experiments suggest that this neuromodulator reinforces stability in idiosyncratic individual behavior. Further study is necessary to determine if serotonin regulates drift through its role in synaptic plasticity or some other widespread effect on the *Drosophila melanogaster* nervous system.

The combination of genetic and neuromodulatory control suggests that extent of behavioral drift could evolve if it confers a fitness advantage in natural settings. The bet-hedging framework provides a theoretical basis for the adaptive value of stable idiosyncrasy in behavior(2, 49): namely, that variability in progeny phenotypes increases the chances that at least some offspring are fit when the environment fluctuates unpredictably. We hypothesize that behavioral drift may be advantageous for similar reasons, but operating within the lifespan of each individual. We provide two arguments in support of this hypothesis. First, we show in several analytically tractable cases (Fig 2A, Supplemental Materials) that it is evolutionarily optimal for an individual to switch behavioral preference phenotypes with a probability equal to the probability of the environment changing to favor those behavioral phenotypes.

This analytical result matches what we find using a more biologically realistic computational model of drifting individual preferences in fluctuating environments. We find that phenotypic drift is beneficial when the environment changes faster than the time to reproductive maturity of an animal. This finding suggests that behaviors that confer fitness with respect to aspects of the environment that change quickly should correspondingly change more quickly. Conversely, stable behavioral preferences may confer fitness with respect to stable aspects of the environment. This is borne out by simulating populations in different environments: quickly changing environmental parameters such as relative humidity (Figures 4A, S4D) favor a drift strategy, while environments that vary at longer (e.g., seasonal) time scales, such as average daily temperature, favor a bet-hedging strategy (Figures 4B, S4D). A key parameter of this model is the age of reproductive maturity (*a*_min_). Phenotypic drift provides a mechanism for organisms with slow development times to survive fluctuating environments long enough to reproduce. Thus, a key quantity is the time scale of environmental fluctuations relative to maturation time. We predict that organisms with more delayed onsets of reproduction will be more likely to exhibit phenotypic drift compared to organisms with more rapid development (subject to the same environmental fluctuations).

Organisms employ many strategies to survive changing environments. While our study focused on the respective advantages of bet-hedging (stable variability at the individual level) and phenotypic drift (variability within an individual’s lifespan), these findings complement previous work comparing bet-hedging and adaptive tracking (selection-induced changes in allelic frequency over time(50)). Previous work on thermal preference in flies suggested that adaptive tracking offers an advantage when the environment fluctuates on approximately years-long time scales, whereas bet-hedging offers an advantage for months-long fluctuations (6, 30). These time scales roughly correspond to several fly lifespans and one lifespan, respectively. This study finds that phenotypic drift may be an adaptive strategy for environmental fluctuations within a lifespan. Thus, adaptive tracking, bet-hedging and phenotypic drift appear to be complementary strategies for the challenges of environmental fluctuation over a wide range of time scales.

Importantly, random phenotypic drift and bet-hedging come at potentially high costs. If organisms could detect (or predict) environmental fluctuations and deterministically change their phenotype to be optimal for the realized environment, they could avoid the losses of randomly choosing the wrong phenotypes. Nonetheless, drift may play an adaptive role in some systems for two key reasons. Firstly, sensing and predicting future environments may be metabolically costly (19), especially for all possible environments. Random changes in preferences may therefore be more efficient than evolving the ability to reliably adapt to any fluctuation.

Second, random strategies may be inherently optimal. In a game theoretic context, finite games always have an optimal strategy (a Nash equilibrium), but this strategy may have to be random. For instance playing randomly in Rock Paper Scissors is optimal in so far as your opponent cannot learn to predict your play and exploit it. The randomness of stochastic evolutionary strategies, such as bet-hedging and phenotypic drift, may thus be truly optimal(41, 42). This may be particularly likely if the fitness effect of a particular behavioral phenotype depends on fluctuating game-theoretic interactions.

The interaction between behavioral drift and learning is likely complicated. Studies have observed idiosyncratic differences in learning that help shape the population-level distribution (51, 52). Specifically, variability in response to identical cues may lead to variation in the population. Similarly, non-adaptive plasticity in response to unrelated cues has been shown to be a mechanism for generating differences in a population as part of a bet-hedging strategy (53). While we limit our analysis in this paper to random shifts in phenotype rather than adaptive plasticity and learning, all these mechanisms of change likely co-exist in natural behaviors.

This paper characterizes changes in individual behavior within a fly’s lifetime, and proposes a theoretical framework in which such changes are evolutionarily adaptive. Together, these results motivate continued study of the biological mechanisms underpinning behavioral drift as well as empirical studies to test the hypothesis that behavioral drift helps organisms survive rapidly changing environments.

## Materials and Methods

See SI Materials and Methods for details. All raw data, Data Acquisition Software, and analysis scripts are available at http://lab.debivort.org/drift-in-individual-preference/ and https://zenodo.org/doi/10.5281/zenodo.13698148. Analysis scripts are available at https://github.com/Maloney-Lab/Drift-in-Individual-Preference.git

### Fly Care

Flies were raised at room temperature and maintained at room temperature. Flies were fed cornmeal/dextrose medium, as previously described (5). Flies were maintained in communal housing before the first time point (Age 2-5), and then maintained in individual housing to maintain identity at later timepoints. Flies were kept on normal 12:12h light dark cycles.

### Continuous Circling Experiments

For the 24 hr continuous circling experiments (Fig 1 A-C, Fig S1A-D), flies were placed in a 28mm radius circular arena filled with standard fly food allowing 3mm of space between food floor and lid. Arenas were illuminated with white light from 9am to 9pm. Flies were tracked with MARGO, as described previously (54).

### Handedness

For Y-maze experiments, flies were placed in Y-mazes as described previously (3, 54). Flies were allowed to roam freely in the Y-maze for two hours, and the number of left and right turns were computed for each fly. Y-mazes were illuminated in white light and flies were tracked with MARGO(54).

### Statistics and Modeling

Data were analyzed in Matlab and Python. Bayesian analysis was performed using the STAN programming language (55). All simulations were done in Python.

### Real-world Data

Data was collected from publicly available sources (NEON(44) and NOAA(43)) for 43 environmental variables across 118,618 sites sampled from the last 20 years.

## ACKNOWLEDGMENTS

We thank M. Miyagi, S. Lavopulo, S. Lall, D. Lavrentovich, and other members of the de Bivort lab for helpful comments on this manuscript. This work was funded by NIH R01NS121874-01 to BdB and a Harvard Brain Institute Postdoctoral Pioneers grant to RM.

