## Supplemental Material for "Drift in Individual Behavioral Phenotype as a Strategy for Unpredictable Worlds"

#### This PDF file includes:

Supporting text
Figs. S1 to S4
Tables S1 to S3
SI References

### Supporting Information Text

### 1. Proof of optimal rate of switching phenotypes for two-state model

Consider a population facing the possibility of a shift in the environment with a probability $p_{\text{shift}}$. Each individual can choose to adopt a behavioral phenotype a new behavioral phenotype or stick with its current phenotype. If the phenotype that it chooses does not match the forthcoming environment, it will die. Otherwise, it will survive and reproduce with rate $\beta$. Evolution will maximize the long-term geometric growth rate.

**Proposition 1.1.** *In the case of two environments, the optimal fraction of progeny to shift to a phenotype adaptive to the different environment ($f_{shift}$) is equal to the probability of the environment changing to the other state ($p_{shift}$).*

*Proof.* The total change in population $G$, conditional on whether the environment shifts or stays, depends on the fraction $f$ of animals that shift or stay in their behavioral phenotype.

$$G_{\text{shift}} = \beta f_{\text{shift}},$$
$$G_{\text{stay}} = \beta f_{\text{stay}}.$$

Since population growth is multiplicative, the long-term change in population over $T$ generations, of which $T_{\text{shift}}$ have a change in the environment and $T_{\text{stay}}$ have no change is given by:

$$G_{\text{total}} = G_{\text{shift}}^{T_{\text{shift}}} G_{\text{stay}}^{T_{\text{stay}}},$$

and the long-term geometric average growth rate as

$$G_{\text{avg}} = (G_{\text{shift}}^{T_{\text{shift}}} G_{\text{stay}}^{T_{\text{stay}}})^{\frac{1}{T}}.$$

In the limit of large $T$,

$$T_{\text{shift}} = T p_{\text{shift}},$$
$$T_{\text{stay}} = T(1 - p_{\text{shift}}).$$

We can then write $G_{\text{avg}}$ in terms of $f_{\text{shift}}$ and $p_{\text{shift}}$:

$$G_{\text{avg}} = \beta f_{\text{shift}}^{p_{\text{shift}}} \cdot \beta(1 - f_{\text{shift}})^{1-p_{\text{shift}}},$$

and the logarithm of average growth becomes

$$\log G_{\text{avg}} = 2\log\beta + p_{\text{shift}} \log f_{\text{shift}} + (1 - p_{\text{shift}}) \log(1 - f_{\text{shift}}).$$

We can find the value of $f_{\text{shift}}$ where $G_{\text{avg}}$ is maximized by taking the partial derivative of $G_{\text{avg}}$ with respect to $f_{\text{shift}}$ to get

$$\frac{\partial \log(G_{\text{avg}})}{\partial f_{\text{shift}}} = \frac{(f_{\text{shift}} - p_{\text{shift}})}{f_{\text{shift}}(f_{\text{shift}} - 1)},$$

which equals zero when $f_{\text{shift}} = p_{\text{shift}}$. As $0 < f_{\text{shift}} < 1$, the second derivative is negative for all values of $f_{\text{shift}}$ and $p_{\text{shift}}$ thus $f_{\text{shift}} = p_{\text{shift}}$ maximizes $G_{\text{shift}}$. □

### 2. Proof of optimal distribution of phenotypes for *n* Discrete states

Consider the discrete-time stochastic process $\{e_t\}$ as a simple model of some fluctuating environmental signal. For the moment, we assume a finite set of possible environmental states. Further, we suppose that the process is strictly stationary, so that all $e_t$ can be regarded as i.i.d. draws from the same categorical distribution $(p_e)$ over environmental states $e \in \{1, 2, \ldots, m\}$.

Suppose that an organism expresses exactly one phenotype from birth, according to the categorical distribution $(f_\phi)$ over phenotypes $\phi \in \{1, 2, \ldots, n\}$. Let $\beta_{e_t,\phi}$ be the fitness of phenotype $\phi$ under environmental state $e$, i.e., the subpopulation expressing phenotype $\phi$ will grow by a factor of $\beta_{e_t,\phi}$ during generation $t$. After $k$ generations (and assuming that the population size is so large as to be infinitely divisible), the total population will have changed by a factor of

$$G_k = \prod_{t=1}^{k} \sum_{\phi=1}^{n} \beta_{e_t,\phi} f_\phi, \qquad e_t \sim \text{Categorical}(p_1, \ldots, p_m). \tag{1}$$

**A. Diagonal fitness.** Suppose that the fitness matrix $\beta = [\beta_{e\phi}]$ is diagonal ($\beta_{e,\phi} = \delta_{e,\phi}\beta_\phi$), and, for simplicity, square ($m = n$), so that only phenotype $\phi = e$ has nonzero fitness in a given environment $e$. We then have

$$G_k = \prod_{t=1}^{k} \beta_{e_t} f_{e_t}, \qquad e_t \sim \text{Categorical}(p_1, \ldots, p_n). \quad [2]$$

Kelly(1) made the following crucial observation:

**Proposition 2.1** (Kelly)**.** *The phenotype distribution* $(f_e)$ *that maximizes the expected population fold change* $\langle G_1 \rangle$ *over a single generation is simply* $f_e = \delta_{e\hat{e}}$*, where* $\hat{e} = \operatorname*{argmax}_e p_e$*.*

In other words, the strategy that maximizes the expected population increase over any finite time horizon is for *all* individuals to express the unique phenotype adapted to the maximally likely environment. This strategy will result in certain extinction as soon as a different environment occurs, which will happen almost surely in the limit that $k \to \infty$. For this reason, Kelly introduced an alternative measure of population fitness: namely, the time-averaged exponent of growth.(1).

**Definition 2.1.** We define the *asymptotic growth rate* to be

$$\lambda = \lim_{k \to \infty} \frac{1}{k} \log G_k, \quad [3]$$

and we say that any phenotype distribution $(f_e)$ that maximizes $\lambda$ is *Kelly-optimal*.

The following theorem due to Bregman (2) justifies the use of the Kelly criterion.

**Theorem 2.1** (Bregman)**.** *A Kelly-optimal phenotype distribution is superior to all other betting strategies in two senses:*

1. *The growth rate of a Kelly-optimal strategy asymptotically dominates that of any other strategy almost surely.*
2. *A Kelly-optimal strategy minimizes the expected time for the population size to increase by an arbitrarily large amount.*

**Proposition 2.2.** *The Kelly-optimal phenotype distribution for the case of a diagonal fitness matrix is* $f_e = p_e$*, independent of* $\beta$*.*

*Proof.* Using the law of large numbers, we can replace the time average in Eq. (3) by an expected value:

$$\begin{aligned}
\lambda &= \lim_{k \to \infty} \frac{1}{k} \log G_k && [4]\\
&= \lim_{k \to \infty} \frac{1}{k} \sum_{t=1}^{k} \log\left(\beta_{e_t} f_{e_t}\right) && [5]\\
&= \sum_{e=1}^{n} p_e \log\left(\beta_e f_e\right). && [6]
\end{aligned}$$

Which can be rearranged as follows:

$$\begin{aligned}
\lambda &= \sum_{e=1}^{n} p_e \log \beta_e + \sum_{e=1}^{n} p_e \log f_e \frac{p_e}{p_e} && [7]\\
&= \underbrace{\sum_{e=1}^{n} p_e \log \beta_e}_{\langle \beta_p \rangle} + \underbrace{\sum_{e=1}^{n} p_e \log p_e}_{-H(p)} - \underbrace{\sum_{e=1}^{n} p_e \log \frac{p_e}{f_e}}_{D_{\mathrm{KL}}(p \| f)}. && [8]
\end{aligned}$$

These three terms are the expected value of $\beta_p$, the negative entropy of $p$, and the Kullback–Leibler divergence of $p$ from $f$, respectively. Only the third term depends on the phenotype distribution $f$. Since $D_{\mathrm{KL}}(p \parallel f) \geq 0$, with equality if and only if $f_e = p_e$, $\lambda$ is maximized when $f_e = p_e$. □

The expression for $\lambda$ given in Eq. (8) has a very intuitive interpretation. To see this, we note that the first term is simply the asymptotic growth rate for a population whose members *all* perfectly match the environment at every generation. However, a population will in general do worse than this upper bound for two reasons: (1) the intrinsic unpredictability of the environment (reflected by a penalty of magnitude $H(p)$, the entropy of the environment), and (2) incomplete knowledge of the true environmental distribution (reflected by a penalty of magnitude $D_{\mathrm{KL}}(p \parallel f)$, the K–L divergence between the phenotype and environment distributions).

Yet another way of understanding the intuition behind Proposition 2.2 is motivated by considering the *worst-case* value of $\lambda$ under the Kelly-optimal phenotype distribution: i.e., by minimizing $\lambda_{\text{Kelly}} = \langle\beta\rangle_p - H(p)$ w.r.t. the environment distribution $(p_e)$. Using a Lagrange multiplier to enforce the normalization of $p$, one obtains

$$p_e^* = \underset{\|p\|=1}{\operatorname{argmin}}\, \lambda_{\text{Kelly}} = \frac{1}{z\beta_e}, \quad [9]$$

where $z = \sum_e \beta_e^{-1}$ is a normalizing constant. Evidently the asymptotic growth rate is minimized when the probability of a given environment is inversely proportional to the fitness of the matching phenotype, so that the organism thrives only in low-probability environments. Using Eq. (9), we may then rewrite Eq. (8) in the form

$$\lambda = -\log z + \underbrace{\sum_{e=1}^{n} p_e \log p_e z\beta_e}_{D_{\text{KL}}(p\|p^*)} - \underbrace{\sum_{e=1}^{n} p_e \log \frac{p_e}{f_e}}_{D_{\text{KL}}(p\|f)}. \quad [10]$$

Thus, the fitness achieved by an organism reflects two opposing K-L divergences corresponding to the how far the environmental distribution is from one that minimizes the Kelly-optimal growth rate ($D_{\text{KL}}(p \parallel p^*)$) and how far the environmental distribution is from the organisms phenotypic distribution ($D_{\text{KL}}(p \parallel f)$). By analogy, fitness depends on the respective degrees to which an organism and its "antagonists" (i.e., the forces that conspire to determine the fitness landscape $\beta$) are maximally informed about the distribution of environmental states. In the case that $z = 1$, the overall growth rate is positive exactly when the organism "knows" more about the environment than the antagonists, in the sense that the environment distribution more closely matches the organism's phenotype distribution than it does the inverse fitness landscape. Moreover, we may regard the term $-\log z$ as a measure of "excess fitness," or the degree to which the environment overall is biased in favor of the organism: if it is negative ($z > 1$), then the organism has to do *better* than the antagonists just to break even (i.e., to achieve $\lambda = 0$), while if it is positive ($z < 1$), then the organism can still achieve net positive growth even under the worst-case scenario that $p_e \propto \beta_e^{-1}$.

**B. Constant baseline fitness.** Suppose that $\beta_{e,\phi} = \epsilon + \delta_{e,\phi}(\tilde\beta_\phi - \epsilon)$, corresponding to the case of a fitness matrix of the form

$$[\beta_{e,\phi}] = \begin{pmatrix} \tilde\beta_1 & \epsilon & \epsilon & \cdots & \epsilon \\ \epsilon & \tilde\beta_2 & \epsilon & \cdots & \epsilon \\ \epsilon & \epsilon & \tilde\beta_3 & \cdots & \epsilon \\ \vdots & \vdots & \vdots & \ddots & \vdots \\ \epsilon & \epsilon & \epsilon & \cdots & \tilde\beta_n \end{pmatrix},$$

where $\epsilon > 0$ is the (small) baseline fitness in the case of a phenotype–environment mismatch. The asymptotic growth rate is then given by

$$\lambda = \lim_{k\to\infty} \frac{1}{k} \log G_k \quad [11]$$

$$= \lim_{k\to\infty} \frac{1}{k} \sum_{t=1}^{k} \log\left(\sum_{\phi=1}^{n} \beta_{e_t,\phi} f_\phi\right) \quad [12]$$

$$= \sum_{e=1}^{n} p_e \log\left(\sum_{\phi=1}^{n} \beta_{e,\phi} f_\phi\right) \quad [13]$$

$$= \sum_{e=1}^{n} p_e \log\left(\tilde\beta_e f_e + \epsilon(1 - f_e)\right). \quad [14]$$

**Claim 2.1.** *To leading order in $\epsilon$, the Kelly-optimal phenotype distribution in the case of constant baseline fitness is given by*

$$f_e \simeq p_e(1 + \epsilon z) - \frac{\epsilon}{\tilde\beta_e},$$

*where $z = \sum_e \tilde\beta_e^{-1}$ as above.*

*Proof.* We seek a first-order solution to the following constrained optimization problem:

$$\min_f \quad \sum_{e=1}^{n} p_e \log\left(\tilde\beta_e f_e + \epsilon(1 - f_e)\right) \quad [15]$$

$$\text{s.t.} \quad \sum_{e=1}^{n} f_e = 1 \quad [16]$$

Applying the method of Lagrange multipliers to the Lagrangian function

$$\mathcal{L}(f,\mu) = \sum_{e=1}^{n} p_e \log\left(\tilde{\beta}_e f_e + \epsilon(1-f_e)\right) + \mu\left(1 - \sum_{e=1}^{n} f_e\right), \quad [17]$$

we obtain the following set of equations:

$$0 = \frac{\partial \mathcal{L}}{\partial f_e} = \frac{p_e(\tilde{\beta}_e - \epsilon)}{\tilde{\beta}_e f_e + \epsilon(1-f_e)} - \mu, \qquad e = 1..n, \quad [18]$$

$$0 = \frac{\partial \mathcal{L}}{\partial \mu} = 1 - \sum_{e=1}^{n} f_e. \quad [19]$$

Linearizing Eq. (18) yields

$$\mu = \frac{p_e}{f_e}\left(1 - \frac{\epsilon}{\tilde{\beta} f_e} + O(\epsilon^2)\right), \quad [20]$$

which we may solve to obtain

$$f_e = \frac{p_e \pm p_e}{2\mu} \mp \frac{\epsilon}{\tilde{\beta}_e} + O(\epsilon^2). \quad [21]$$

We reject the solution $f_e = \epsilon/\tilde{\beta}_e$ on grounds of non-normalizability. Plugging into Eq. (19) then yields

$$1 = \sum_{e=1}^{n} f_e = \frac{1}{\mu} - \epsilon \underbrace{\sum_{e} \frac{1}{\tilde{\beta}_e}}_{z} \implies \mu = \frac{1}{1+\epsilon z}, \quad [22]$$

from which it follows that

$$f_e = p_e(1+\epsilon z) - \frac{\epsilon}{\tilde{\beta}_e} + O(\epsilon^2), \quad [23]$$

as desired. □

As expected, this asymptotic solution reduces to that of Proposition 2.2 in the limit of $\epsilon \to 0$. However, at finite $\epsilon$, we find that the optimal probability allocated to phenotype $e$ is now reduced by an amount inversely proportional to that phenotype's on-diagonal fitness $\tilde{\beta}_e$. Thus, unlike in the fully diagonal case, the optimal phenotype distribution $(f_e)$ under an assumption of nonzero baseline fitness depends on the precise fitness landscape. In particular, relatively low-fitness phenotypes should now occur *less* frequently than their matching environments. The corresponding Kelly-optimal growth rate is

$$\lambda \simeq \tilde{\lambda} + \epsilon\left(z - \sum_{e=1}^{n} \frac{p_e}{\tilde{\beta}_e}\right) \quad [24]$$

$$= \tilde{\lambda} + \epsilon \sum_{e} \frac{1-p_e}{\tilde{\beta}_e}, \quad [25]$$

where $\tilde{\lambda}$ is the optimal growth rate Eq. (10) in the case of purely diagonal fitness.

### 3. A continuum of states

We consider the case in which the state space $E$ is uncountably infinite, indexed by a single scalar $e$, so that the environment $e_t$ becomes a real-valued (though still discrete-time and stationary) stochastic process. We assume that the phenotype space is continuous and isomorphic to $E$, such that it, too, can be indexed by $e$. Then the categorical distributions $(p_e)$ and $(f_\phi)$ of environments and phenotypes, respectively, become density functions $p(e)$ and $f(\phi)$. Likewise, the fitness matrix $\beta_{e\phi}$ becomes a continuous function $\beta(e,\phi)$ whose bandwidth controls the "sharpness" or "fuzziness" of the correspondence between environmental states and phenotypes. By analogy to Eq. (1), we have

$$G_k = \prod_{t=1}^{k} \int_E \beta(e_t,\phi) f(\phi) \mathrm{d}\phi, \quad [26]$$

where $e_t$ are i.i.d. samples from the distribution with density $p(e)$. One possible form of $\beta(e,\phi)$ is proportional to a Gaussian with standard deviation $\sigma$ centered at $\phi = e$:

$$\beta(e,\phi) = \frac{\hat{\beta}(e,\phi)}{\sqrt{2\pi\sigma^2}} e^{-\frac{(\phi-e)^2}{2\sigma^2}}. \quad [27]$$

When $\sigma$ is small, we may use Laplace's method (equivalent to considering the limit $\sigma \to 0$) to write

$$G_k = \prod_{t=1}^{k} \int_E \beta(e_t, \phi) f(\phi) \mathrm{d}\phi \quad [28]$$

$$= \prod_{t=1}^{k} \int_E \frac{\hat{\beta}(e_t, \phi) f(\phi)}{\sqrt{2\pi\sigma^2}} e^{-\frac{(\phi - e_t)^2}{2\sigma^2}} \mathrm{d}\phi \quad [29]$$

$$= \prod_{t=1}^{k} \left[\hat{\beta}(e_t, e_t) f(e_t) + O(\sigma^2)\right]. \quad [30]$$

We have thus reduced the problem to the discrete case of Eq. (2) treated above, so that the analogue of Proposition 2.2 follows trivially.

**Claim 3.1.** *For $\sigma$ small, the phenotype distribution $f(\phi)$ that maximizes the asymptotic growth rate $\lambda$ is just $f(\phi) \simeq p(e)$, independent of the fitness landscape $\beta(e, \phi)$.*

### Supplementary Methods

**Data and Code.** All data described in this paper and analysis code are available at http://lab.debivort.org/drift-in-individual-preference/ and at https://zenodo.org/doi/10.5281/zenodo.13698148 and https://github.com/Maloney-Lab/Drift-in-Individual-Preference.git (code only).

**Fly Care.** Flies were grown on standard cornmeal/dextrose medium as previously described (3). Flies were kept on 12:12h light:dark cycle at room temperature (20-23°). All behavioral experiments (excepting continual tracking experiments) were done during the animals' subjective day. Unless otherwise mentioned, all flies were 3-6 days post eclosion at the time of their first use in a behavior experiment. Fly stocks used in this experiment are described in Table S1. To maintain individual identity across experiments, flies were either stored alone in standard media vials or in individual housing fly plates (modified 96 well plates (4); FlySorter, LLC).

**Continuous Circling Experiments.** For continuous circling experiments, circular arenas were fabricated from three layers of laser-cut acrylic (floor, wall and lid-holder layers) (5). Lids were cut from 3mm thick clear acrylic). Wall layers were made from black acrylic an defined arenas of radius 28mm and depth of either 1.6mm or 10mm. The latter arenas were filled with standard fly food until 1.6mm of clearance remained. Flies were anaesthetized by $CO_2$ and loaded singly into arenas, allowing 30 minutes of acclimation post anaesthetization. Transparent lids coated in sigmacote (Sigma-Aldrich) were placed on top of arenas to prevent fly escape and ceiling walking. Flies were recorded using a PointGrey Blackfly BFLY-PGE 12A2M camera at 30 frames per second for up to 14 days: either twice daily for 2h per session, or continuously for 24h. In the former experiment, fly identity was maintained between sessions by individual storage as described above. In the continuous experiment, flies were transferred to arenas with fresh food after anesthetization with $CO_2$.In 2 hr experiments, flies were given 30 minutes to recover before experiments. We used MARGO (6) to record fly centroids in real time. All assays were conducted under 12:12h light and dark cycle conditions (9AM:9PM).

Power spectra were calculated using Lomb-Scargle periodograms (7) to accommodate gaps in data due to lack of fly activity and periods when behavior was not recorded. Daily low pass filtered data (Fig 1B) were creating using a Blackman filter (8) with a window size of 51 h, omitting missing data.

**Y-Maze Experiments.** Y maze experiments were performed as described previously (6, 9). Briefly, flies were loaded individually into symmetrical y-maze arenas and their centroid locations were tracked using MARGO (6), which detected each turn (moving from one arm of the y-maze to another) and calculated the fraction of right turns. All experiments were performed in white LED light in order to increase the activity of flies during the flies subjective day photoperiod. Flies were anesthetized with $CO_2$ prior to loading into arenas, flies were allowed to acclimate for 20 minutes in the arenas prior to the beginning of the experiment. Handedness was computed based on the turns made in a 2 hour period each day, turns were scored based on the decision made each time the fly entered the center of the y-maze. Flies were cold anesthetized for removal from the arenas and identity was maintained as described above.

**Bayesian Inference.** Estimates for $\sigma_{\text{bet-hedging}}$, $\sigma_{\text{drift}}$, and $\varphi$ were generated using MCMC sampling in STAN 2.35 (10) according to the model in Fig 1F. Weakly informative priors were used as follows: $\sigma_{\text{bet-hedging}}$=Inverse Gamma($\alpha$=3, $\beta$=1); $\sigma_{\text{drift}}$=Inverse Gamma($\alpha$=3, $\beta$=1); $\varphi$=Normal($\mu$=0, $\sigma^2$=10) $R_{\text{Missing}}$=Normal($\mu$=0, $\sigma^2$=10). MCMC was performed with 4 chains, with the first 1000 draws discarded, posteriors were determined based on the subsequent 2000 draws. Inferred posteriors were robust to the choice of weakly informative priors.

Q values in 1H,I were estimated based on the minimum posterior probability of the measured parameter having a difference of zero or less relative to the observed relationships based on the MCMC draws in STAN. In the case where the full simulated distribution had no overlap (Fig 1H, Phi), q values were given as q<1e-7.

**Pharmacology.** For experiments where flies were treated with aMW, flies were collected as pupae and allowed to eclose on either treated or untreated food. Food was prepared and concentrations chosen based on previous reports (11, 12). For the aMW condition, aMW was added a a concentration of 20 mM with ascorbic acid used as a stabilizer (25 mg of ascorbic acid for every 100 mL of fly food). For the 5-HTP condition, 5-HTP was added at a concentration of 50mM. Experiments (and corresponding control flies) were limited to flies who were at least 4 days post-eclosion at the start of the first timepoint. Food was replaced weekly.

**Generation of $trh^n$ mutant.** To generate a $trh^n$ mutant in a defined genetic background, we used *in vivo* CRISPR (13) to create $trh^n$ mutants in the Isod1 background (an isogenized Oregon-R derivative)(14). Briefly, Act-Cas9; Bl/CyO; TM2/TM6b flies were crossed to y/Y;U6-sgRNA flies (BDSC 91886) to produce +/y; +/CyO;$trh^n$/Tm6B flies. Mutations were verified via Sanger sequencing to generate a frame shift mutation at the targeted site. which were crossed back to +(Isod1);+(Isod1);Tm2/TM6b flies for multiple generations to produce +(Isod1);+(Isod1);$trh^n$ flies, where all chromosomes except for the third chromosome (with the null mutant trh) were identical to IsoD1 controls. Putative mutant lines were genotyped via sanger sequencing, $trh^n$ lines used in this study were selected based on verification that non-homologous end joining induced a frame-shift mutation at the targeted point.

**Simulations.** Simulations were implemented in Python 3 based on the model described in Fig 2a. All scripts are available at http://lab.debivort.org/drift-in-individual-preference/ and at https://zenodo.org/doi/10.5281/zenodo.13698148 and https://github.com/Maloney-Lab/Drift-in-Individual-Preference.git. Changes in population were evaluated at steps corresponding to one day, and continuous distributions of preference were approximated with 200 preference bins ranging from an arbitrary scaling of -1 to 1. Variations in amount of daily drift ($\sigma_{\mathrm{drift}}$), dispersion of initial preferences ($\sigma_{\mathrm{drift}}$), the bounding envelope($\sigma_{\mathrm{max}}$), the size of fluctuations in the environment ($\sigma_{\mathrm{mean}}$) and the width of the environmental filter ($\sigma_{\mathrm{max}}$) are all calculated on this $[-1, 1]$ range. Flies were assumed to have a maximum lifespan of ten days plus twice the age of reproductive maturity for computational efficiency. Increasing maximum lifespan past this point did not quantitatively change the results of the simulations. Results for most simulations presented in the manuscript are available at the above URLs. For some results, only the environments are provided for space reasons, as the simulation is deterministic and can be rerun.

Environmental time-series ($E_t$) were created by generating random white noise time series and band-pass filtering them to the specified frequency bands in python using the `np.fft` library. Resulting filtered time series were normalized to have a mean of zero and a standard deviation of $\sigma_{\mathrm{mean}}$.

Detailed parameters of simulations used in figures are shown in Table S2.

**Real-World Data Collection.** Real world time series with one-day (or better) resolution were collected from the Global Historical Climatology Network daily (GHCNd) and National Ecological Observatory Network (NEON) datasets. The GHCNd that is produced by the National Oceanic and Atmospheric Association (NOAA)(15). Timeseries were collected from available stations across the world from 1998 to 2018. We also used data from the National Ecological Observatory Network (NEON) (16). Data was accessed using the neonUtilities package (17) (see Table S2).

Time series used to run simulations was selected based on having at least 1000 days of continuous data with no gaps larger than 5 days: gaps were filled via linear interpolation. Time series with sampling more often than once per day were used to create average, maximum, and minimum daily value time series. Starting dates for sampled time series were chosen randomly among possible dates that met the above criteria such that no data point was used more than once. Due to the heterogeneous nature of the observation sites with respect to time active, available sensors, and data quality, most data types were only available at a subset of sites.

Average Power Spectral Density in Fig S4C was calculated via lomb-scargle and averaged across all available time series.

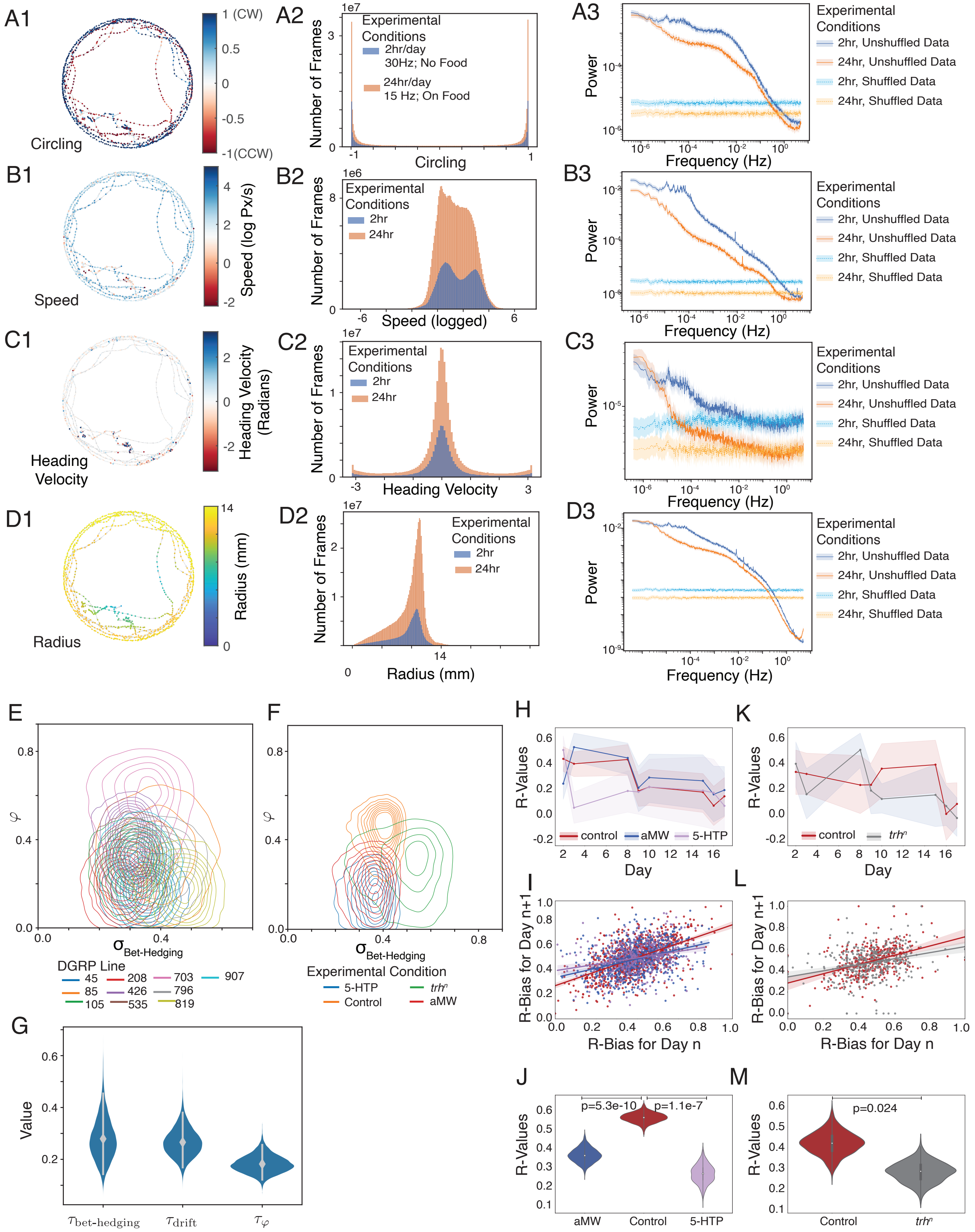


Fig. S1: Change in behavioral biases over time and the influence of genetics and serotonin on rate of change. Caption on next page.

Fig. S1: Change in behavioral biases over time and the influence of genetics and serotonin on rate of change. **A:** Statistics of the "circling" behavior metric. **A1:** Example of one fly's centroid position over time, colored by circling (where -1 equals CCW and 1 equals CW). **A2:** Histogram of circling measurements across all frames and all flies. **A3:** Average power spectral density across all flies as estimated via Lomb-Scargle. Shaded regions indicate bootstrapped 95% confidence intervals (n=1000 replicates). **B1-3:** As A, except for speed (pixels/sec, each pixel=.28mm). **C1-3:** As A-B, except measuring of heading velocity (change in direction between frames of the flies (change in heading angle between timepoints). **D1-3:** As A-C, except measuring the normalized distance of the fly from the center of the arena, in units of unit diameter (28mm) **E:** Bivariate posterior estimates of $\sigma_{\text{Bet-hedging}}$, and $\varphi$ for each DGRP line. Concentric circles represent deciles of posterior density, with the outer circle representing 95% posterior density. **F:** As E, except for experiments with manipulation of serotonin with 5-HTP, AMW, and $trh^{\text{n}}$. **G:** Posterior estimates of distribution of differences between batches in hierarchical model for bet-hedging, drift, and $\varphi$ (See Figure 1 for model). **I:** Estimates of 1 day R value for all pairs of sequential experiments by treatment condition. **J:** Estimates of distribution of possible R by condition, as estimated by bootstrapping. **K-M:** As (I-J), except for experiments with $trh^{n}$ mutants. p values in J,M Fisher Z-Transform

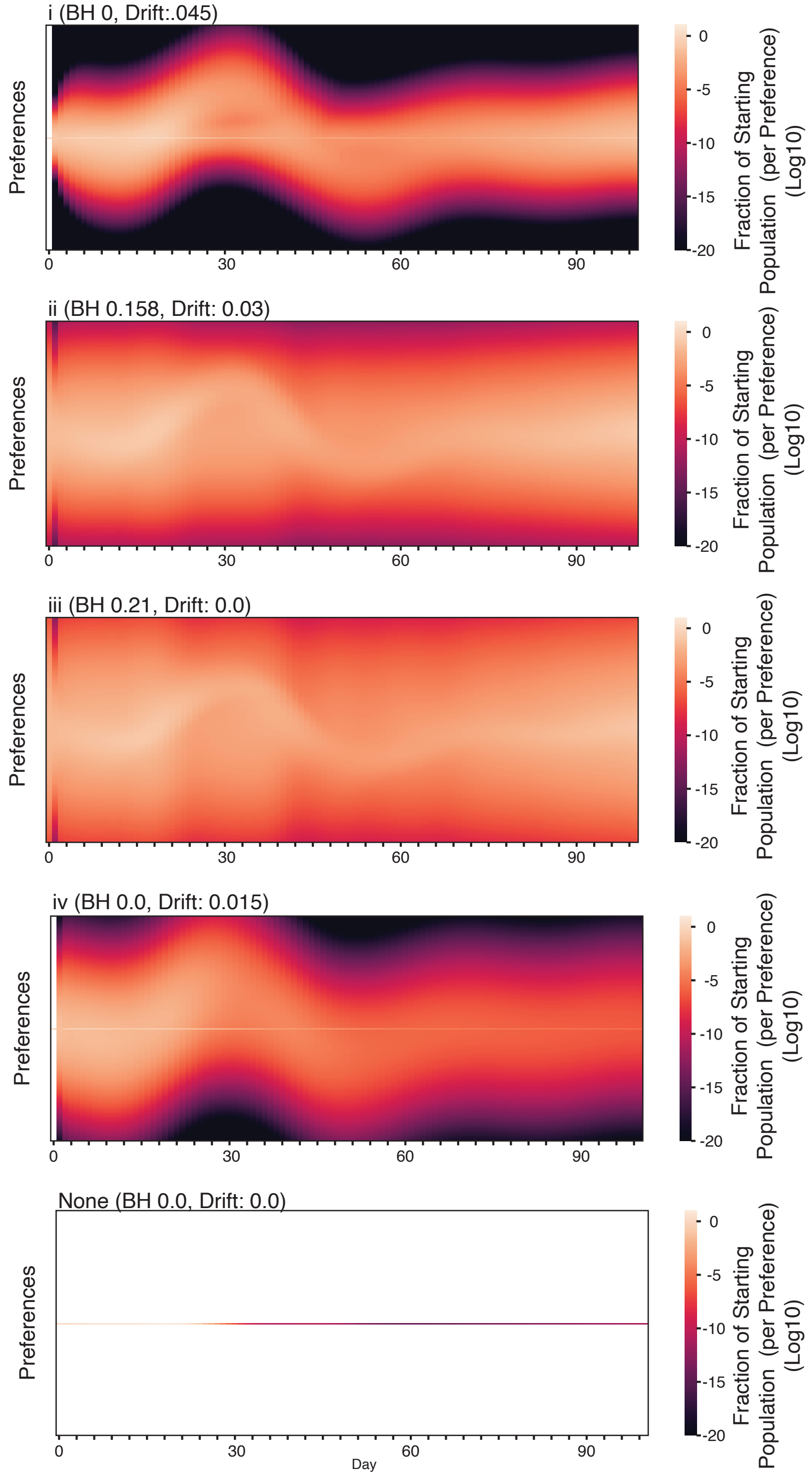


Fig. S2: Distribution of animal preferences over time for different strategies. Strategies from Figure 2C-D, with the fraction of the starting population (colormap) per preference bin (Y axis) plotted over time (x-axis). Mean preferences are reflected in the central preference (as can be noted in cases with no bet-hedging (i,iv, None), as newly born flies begin with the mean preference. The total population summing across all preferences produces the total population graph in 2D.

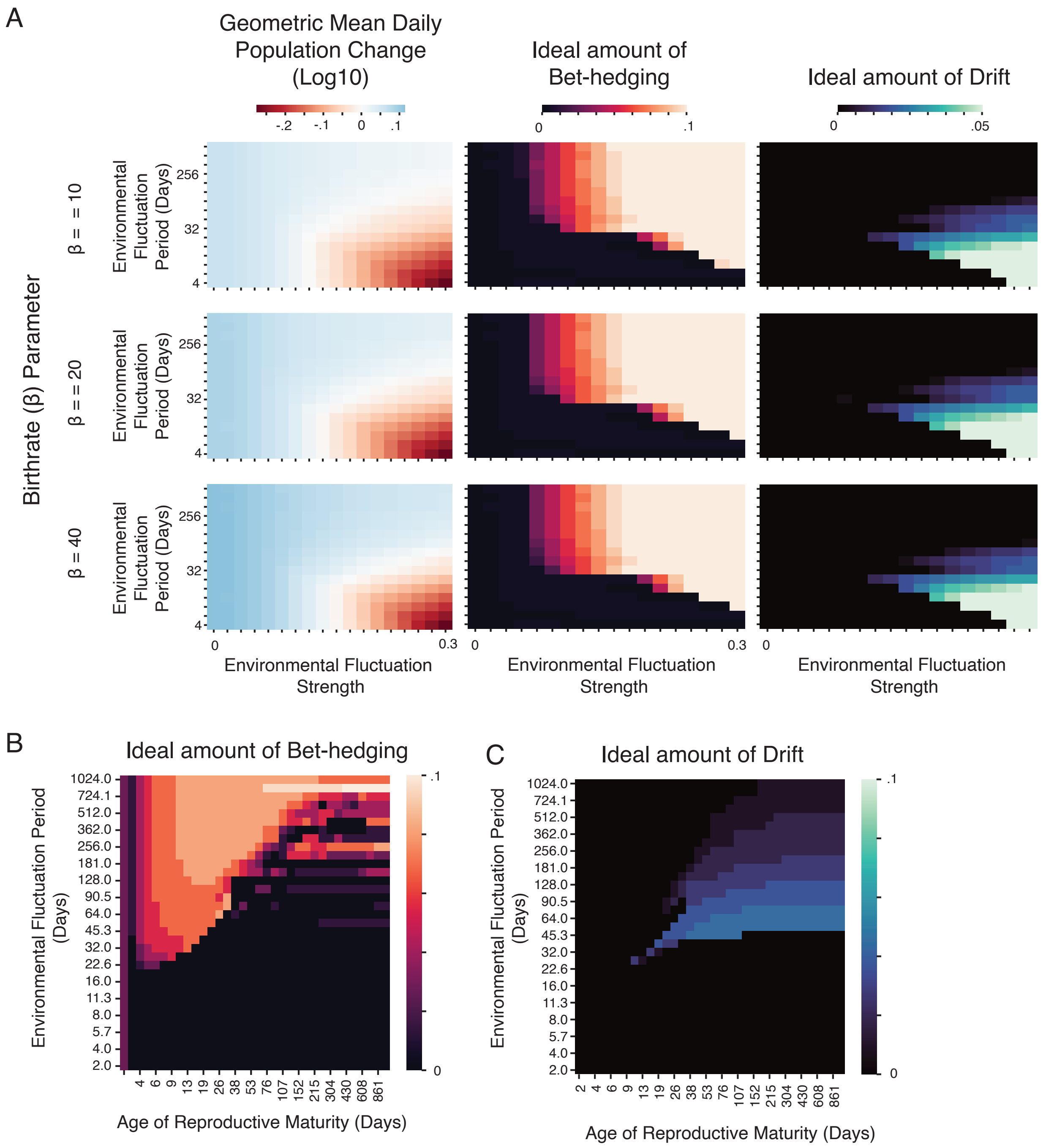


Fig. S3: Effect of birthrate and longer timescales on strategies. **A:** Adjusting birthrate $\beta$ (rows) changes the overall rate of change of the population (left column) without affecting the ideal amount of bet-hedging or drift (middle and right columns). **B:** As Figure 3D, but with logarithmic scale for Fluctuation Period and Age of reproductive maturity.)

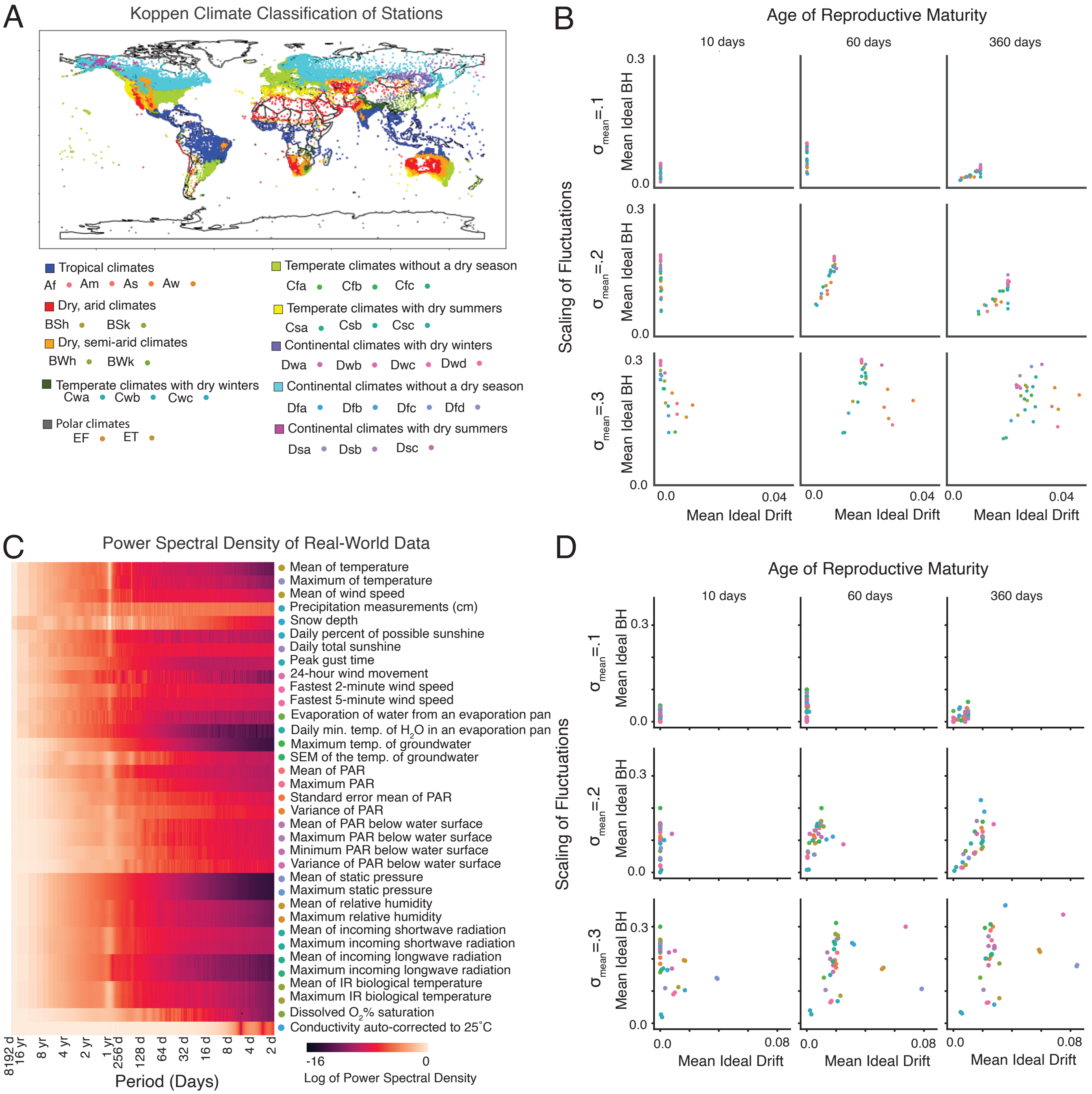


Fig. S4: Breakdown of simulations by Koppen Climate Classification and Measurement Type. **A:** Map of all stations used for the simulation, colored by broad categories of Koppen Climate Clasification. Subcategories indicated with colors for B. **B:** Mean Ideal Drift and Bet-hedging as a function of age of reproductive maturity and scaling of environmental amplitudes, averaged across all time series of a given Koppen climate classification type. **C:** Average Power Spectral density of each environmental measurement type. Colored dots correspond to data in (D). **D:** As (B), but averaged across each type of measurement.

**Table S3. List of NEON Datasets Used**

| Product ID/ DOI/ Data Product Name | Site IDs | Dates | Date Accessed |
|---|---|---|---|
| DP1.20217.001<br>doi.org/10.48443/br51-rd19<br>Temperature of groundwater | WLOU, WALK, TOOK, TOMB, SYCA, SUGG, REDB, PRPO, PRLA, PRIN, POSE, OKSR, MCDI, MAYF, MART, LIRO, LEWI, KING, HOPB, GUIL, FLNT, CRAM, COMO, CARI, BLWA, BLUE, BLDE, BIGC, BARC, ARIK | 2016-03-04 to 2022-04-30 | 06/28/2022 |
| DP1.00024.001<br>doi.org/10.48443/51ss-fm81<br>Photosynthetically active radiation (PAR) | WLOU, WALK, TOOK, TOMB, TECR, SYCA, SUGG, REDB, PRPO, PRLA, PRIN, POSE, OKSR, MCRA, MCDI, MAYF, MART, LIRO, LEWI, LECO, KING, HOPB, GUIL, FLNT, CUPE, CRAM, COMO, CARI, BLWA, BLUE, BLDE, BIGC, BARC, ARIK | 2016-03-04 to 2022-03-31 | 05/04/2022 |
| DP1.20261.001<br>doi.org/10.48443/jnwy-xy08<br>Photosynthetically active radiation below water surface | TOOK, TOMB, SUGG, PRPO, PRLA, LIRO, FLNT, CRAM, BLWA, BARC | 2017-07-28 to 2022-03-31 | 05/04/2022 |
| DP1.00004.001<br>doi.org/10.48443/rt4v-kz04<br>Barometric pressure | YELL, WREF, WOOD, WLOU, WALK, UNDE, UKFS, TREE, TOOL, TOOK, TECR, TEAK, TALL, SYCA, SUGG, STER, STEI, SRER, SOAP, SJER, SERC, SCBI, RMNP, REDB, PUUM, PRPO, PRLA, PRIN, POSE, OSBS, ORNL, ONAQ, OKSR, OAES, NOGP, NIWO, MOAB, MLBS, MCRA, MCDI, MAYF, MART, LIRO, LEWI, LENO, LECO, LAJA, KONZ, KONA, KING, JORN, JERC, HOPB, HEAL, HARV, GUIL, GUAN, GRSM, FLNT, DSNY, DELA, DEJU, DCFS, CUPE, CRAM, CPER, COMO, CLBJ, CARI, BONA, BLUE, BLDE, BLAN, BIGC, BART, BARR, BARC, ARIK, ABBY | 2013-09-12 to 2022-01-31 | 03/13/2022 |
| DP1.00098.001<br>doi.org/10.48443/k9vk-5k27<br>Relative Humidity | ABBY, ARIK, BARC, BARR, BART, BIGC, BLAN, BLDE, BLUE, BONA, CARI, CLBJ, COMO, CPER, CRAM, CUPE, DCFS, DEJU, DELA, DSNY, FLNT, GRSM, GUAN, GUIL, HARV, HEAL, HOPB, JERC, JORN, KING, KONA, KONZ, LAJA, LECO, LENO, LEWI, LIRO, MART, MAYF, MCDI, MCRA, MLBS, MOAB, NIWO, NOGP, OAES, OKSR, ONAQ, ORNL, OSBS, POSE, PRIN, PRLA, PRPO, PUUM, REDB, RMNP, SCBI, SERC, SJER, SOAP, SRER, STEI, STER, SUGG, SYCA, TALL, TEAK, TECR, TOOK, TOOL, TREE, UKFS, UNDE, WALK, WLOU, WOOD, WREF, YELL | 2013-09-12 to 2022-02-28 | 03/14/2022 |

Continued on next page

| Product ID/<br>DOI/<br>Data Product Name | Site IDs | Dates | Date Accessed |
|---|---|---|---|
| DP1.00023.001<br><br>doi.org/10.48443/9qpc-5v70<br><br>Shortwave and longwave radiation (net radiometer) | BART, UKFS, TALL, YELL, WREF, WOOD, WLOU, WALK, UNDE, TREE, TOOL, TOOK, TECR, TEAK, SYCA, SUGG, STEI, SRER, SOAP, SJER, STER, SERC, SCBI, RMNP, REDB, PUUM, PRPO, PRLA, PRIN, POSE, OSBS, ORNL, ONAQ, OKSR, OAES, NOGP, NIWO, MOAB, MLBS, MCRA, MCDI, MAYF, MART, LIRO, LEWI, LENO, LECO, LAJA, KONZ, KONA, KING, JORN, JERC, HOPB, HEAL, HARV, GUIL, GUAN, GRSM, FLNT, DSNY, DELA, DEJU, DCFS, CUPE, CRAM, CPER, COMO, CLBJ, CARI, BONA, BLUE, BLDE, BLAN, BIGC, BARR, BARC, ARIK, ABBY | 2013-09-12 to 2022-05-31 | 06/07/2022 |
| DP1.00005.001<br><br>doi.org/10.48443/jqb2-vy96<br><br>IR biological temperature | YELL, WREF, WOOD, UNDE, UKFS, TREE, TOOL, TEAK, TALL, STER, STEI, SRER, SOAP, SJER, SERC, SCBI, RMNP, PUUM, OSBS, ORNL, ONAQ, OAES, NOGP, NIWO, MOAB, MLBS, LENO, LAJA, KONZ, KONA, JORN, JERC, HEAL, HARV, GUAN, GRSM, DSNY, DELA, DEJU, DCFS, CPER, CLBJ, BONA, BLAN, BART, BARR | 2013-09-12 to 2022-01-31 | 03/14/2022 |
| DP1.20288.001<br><br>doi.org/10.48443/t7rj-pk25<br><br>Water quality | TOOK, TOMB, SUGG, PRPO, PRLA, LIRO, FLNT, CRAM, BLWA, BARC, ARIK, MART, WALK, TECR, SYCA, REDB, PRIN, POSE, OKSR, MCRA, MCDI, MAYF, WLOU, LEWI, LECO, KING, HOPB, GUIL, CUPE, COMO, CARI, BLUE, BLDE, BIGC | 2014-01-11 to 2022-03-16 | 05/04/2022 |

Table S1: *Drosophila melanogaster* genotypes used in this paper

| Genotype | Source | Figure | *n* | Citation |
|---|---|---|---|---|
| Canton-S | BDSC 64349 | 1A-C, S1A-D | 250 (24hr) 252 (2hr) | |
| DGRP 45 | BDSC 28128 | 1E, 1G, S1E | 55 | (18) |
| DGRP 85 | BDSC 28274 | 1E, 1G, S1E | 22 | (18) |
| DGRP 105 | BDSC 28139 | 1E, 1G, S1E | 91 | (18) |
| DGRP 208 | BDSC 25174 | 1E, 1G, S1E | 111 | (18) |
| DGRP 426 | BDSC 28196 | 1E, 1G, S1E | 235 | (18) |
| DGRP 535 | BDSC 28208 | 1E, 1G, S1E | 115 | (18) |
| DGRP 703 | BDSC 28218 | 1E, 1G, S1E | 48 | (18) |
| DGRP 796 | BDSC 28233 | 1E, 1G, S1E | 140 | (18) |
| DGRP 819 | BDSC 28242 | 1E, 1G, S1E | 145 | (18) |
| DGRP 907 | BDSC 28262 | 1E, 1G, S1E | 141 | (18) |
| Isod1 (Oregon R) | Clandinin Lab | 1H-I S(H-M) | 192 each(1H: AMW, 5HTP, Control), 98 (1I: Control) | (14) |
| $trh^n$ | this study | 1I, S1 G,K-M | | |
| GS01997 | BDSC 91886 | | | |
| Act-Cas9 | BDSC 54590 | | | |

Table S2: Model Parameters

| Figure | $e_t$ (Environment time series) | $e_t$ scaling | $\sigma_E$ | $\sigma_D$ | $\sigma_B$ | $\sigma_{max}$ | $a_{min}$ | $\beta(BirthRate)$ | Simulation length (days) |
|---|---|---|---|---|---|---|---|---|---|
| Fig 2 C-2; Fig S3 | Temporally Filtered White Noise | .3 | .125 | 0-.3 | 0-.3 | .3 | 10 | 40 | 100 |
| Fig 2 E | Temporally Filtered White Noise | .3 | .125 | 0-.3 | 0-.3 | .3 | 10 | 40 | 100 |
| Fig 2F | Temporally Filtered White Noise | .3 | .125 | 0-.3 | 0-.3 | .3 | 10 | 40 | 100 |
| Fig 3A;3C | Temporally Filtered White Noise | 0-.3) | .125 | 0-.05 | 0-.01 | .3 | 10 | 40 | 1001 |
| Fig 3B;3D | Temporally Filtered White Noise | .3 | .125 | 0-.05 | 0-.01 | .3 | 2-72 | 40 | 1001 |
| Fig 4A | RH, Arikarree River (NEON), | .1, .3 | .125 | 0-.1 | 0-.1 | .3 | 10,60,360 | 40 | 1001 |
| Fig 4B | Avg. Temp, Longreach AU (NOAA) | .1, .3 | .125 | 0-.1 | 0-.1 | .3 | 10,60,360 | 40 | 1001 |
| Fig 4C-G | NOAA/NEON | .2 | .125 | 0-.1 | 0-.5 | .3 | 10 | 40 | 1001 |
| Fig S3A | Temporally Filtered White Noise | .3 | .125 | 0-.1 | 0-.05 | .3 | 10 | 10,20,40 | 1001 |
| Fig S3B-C | Temporally Filtered White Noise | .3 | .125 | 0-.1 | 0-.1 | .3 | 2-1024 | 40 | 1001 |
| Fig S4B;S4D | NOAA/NEON | .1, .2, .3 | .125 | 0-.3 | 0-.3 | .3 | 10,60,360 | 40 | 1001 |
